# Temperature selection drives evolution of function-valued traits in a marine diatom

**DOI:** 10.1101/167817

**Authors:** Daniel R. O’Donnell, Carolyn R. Hamman, Evan C. Johnson, Christopher A. Klausmeier, Elena Litchman

**Author notes:** **Contributions:** DRO designed all assays, designed and maintained the evolution experiment, conducted all statistical analyses and wrote the manuscript. CRH assisted in experimental design and carried out most of the trait assays. ECJ conducted some trait assays and contributed modeling and statistical expertise. CAK constructed the models and edited the manuscript. EL conceived the idea, designed the experiments with DRO, supervised experiments, provided material, personnel and grant support, and edited the manuscript. **Corresponding author:** Daniel R. O’Donnell, Kellogg Biological Station, 3700 E. Gull Lake Dr., Hickory Corners, MI 49060.

## Abstract

Rapid evolution in response to environmental change will likely be a driving force determining the distribution of species and the structure of communities across the biosphere in coming decades. This is especially true of microorganisms, many of which may be able to evolve in step with rising temperatures. An ecologically indispensable group of microorganisms with great potential for rapid thermal adaptation are the phytoplankton, the diverse photosynthetic microbes forming the foundation of most aquatic food webs. We tested the capacity of a globally important phytoplankton species, the marine diatom *Thalassiosira pseudonana*, for rapid evolution in response to temperature. Evolution of replicate populations at 16 and 31°C for 350-450 generations led to significant divergence in several traits associated with *T. pseudonana’*s thermal reaction norm (TRN) for per-capita population growth, as well as in its competitive ability for nitrogen (commonly limiting in marine systems). Of particular interest were evolution of the optimum temperature for growth, the upper critical temperature, and the derivative of the TRN, an indicator of potential tradeoffs resulting from local adaptation to temperature. This study offers a broad examination of the evolution of the thermal reaction norm and how modes of TRN variation may govern a population’s long-term physiological, ecological, and biogeographic response to global climate change.

## Introduction

The dependence of physiological processes on temperature is perhaps the most important underlying factor determining the fitness of organisms across latitudinal and altitudinal gradients, and thus their distributions and abundances across the planet. In light of recent climate warming, many studies have focused on temperature dependence of physiological traits (e.g. photosynthesis, respiration, per-capita population growth) near the upper bounds of organisms’ thermal tolerance ranges (Rowan 2004; Bradford 2013; Listmann *et al.* 2016) and the ecological consequences of temperatures rising beyond those ranges (Rowan 2004; Thomas *et al.* 2012). The focus of many of these studies is increasingly on the potential for rapid evolution of physiological traits in response to temperature change, and how such evolution may mitigate the ecological impacts of climate warming (Hoffmann & Sgrò 2011; Schlüter *et al.* 2014; Listmann *et al.* 2016).

Evolution on ecological timescales may be commonplace (Schoener, 2011) and can ‘rescue’ populations from potentially catastrophic environmental change (Gomulkiewicz and Holt 1995; Bell 2013; Schiffers *et al.* 2013). This is especially true of microorganisms, which can have enormous population sizes, and many of which reproduce on timescales on the order of minutes to days; these attributes offer many opportunities for mutation and subsequent natural selection to lead to rapid evolution in response to environmental change. Despite the high potential of microbes to adapt to changing environmental conditions, little is known about how a prominent aquatic microbial group, phytoplankton, evolves in response to global change stressors, particularly temperature. Phytoplankton are a diverse group of global importance: they are the foundation of most marine and freshwater food webs and are major drivers of global biogeochemical cycles, carrying out ~50% of global carbon fixation (Field *et al.* 1998) and linking nitrogen and phosphorus cycles (Redfield, 1958). Rising temperatures may negatively impact phytoplankton productivity, biomass and species diversity (Boyce *et al.* 2010; Thomas *et al.* 2012). However, given their high rates of reproduction and large population sizes, some phytoplankton may be capable of rapid evolution in response to warming, mitigating some ecological impacts of global climate change (Litchman *et al.* 2012; Listmann *et al.* 2016). For example, evolutionary change in *T*_opt_ and *CT*_max_ in response to ocean warming may reduce heat-induced mortality and poleward migration of phytoplankton populations, allowing some to persist at low latitudes where they might otherwise go regionally extinct (Thomas *et al.* 2012); such change could mitigate some of the potential broad ecological consequences of regional depletion of phytoplankton diversity.

A number of recent evolution experiments have sought to elucidate how marine phytoplankton evolve to cope with environmental change (Reusch & Boyd 2013), though to date most have focused on increased atmospheric *p*CO_2_ and/or ocean acidification (Jin *et al.* 2013; Crawfurd *et al.* 2011; Lohbeck *et al.* 2012). Surprisingly, thermal adaptation has been studied experimentally in only a couple of phytoplankton species to date, only one of which, a coccolithophorid, was a marine species of global importance (Padfield *et al.* 2015; Schlüter *et al.* 2014; Listmann *et al.* 2016). It is therefore difficult to predict if other ecologically important groups, such as diatoms that contribute up to 25% of all global carbon fixation (Nelson *et al.* 1995), would respond in a similar way. Moreover, while we may expect *T*_opt_ and *CT*_max_ to increase after evolving at high temperatures, knowledge of how other important traits may change in response to elevated temperatures is somewhat limited, to date. An important first step toward understanding multi-trait evolutionary responses to warming was an experimental study by Schlüter *et al.* (2014), who found that after one year of experimental adaptation to elevated temperature (26.3 °C), the marine coccolithophore *Emiliania huxleyi* increased its per-capita population growth rate by up to 16% under both ambient and elevated *p*CO_2_; warm-adapted populations also evolved smaller cell diameter and lower particulate organic and inorganic carbon content compared to cold (15 °C)-adapted populations assayed at 26.3 °C under ambient *p*CO_2_. In a follow-up study, Listmann *et al.* (2016) observed that both the optimal temperature for growth (*T*_opt_) and the maximum temperature at which growth stops (referred to as upper critical temperature, *CT*_max_ herein) increased in warm-adapted populations of *E. huxleyi* compared to cold-adapted strains under both ambient and elevated *p*CO_2_.

While many quantitative traits can defined by a single number (e.g. the depth of a finch’s beak), biological rates in ectotherms vary continuously across temperature—they are thus often referred to as “function-valued traits” (Kingsolver *et al.* 2001). Our understanding of how diverse function-valued traits may evolve in different organisms is limited. Various tradeoffs, such as generalist-specialist or resource-allocation tradeoffs, may be important in determining how the shapes of trait functions, including the thermal reaction norm (TRN), evolve (Angilletta et al. 2003). By definition, the trait values derived from TRN and many other function-valued traits are non-independent across environmental (e.g. temperature) gradients (Angilletta et al. 2003); thus, selection at one temperature may alter biological rates (e.g. population growth rate) at every other temperature along the TRN. Selection may change the slope and curvature at every point as well (Kutcherov 2016), especially if some regions of the TRN are more evolutionarily labile than others (Araújo *et al.* 2013). While recent studies showed that *T*_opt_ and *CT*_max_ increase after evolving at higher temperatures (Listmann et al. 2016), we do not know how the whole TRN may evolve in phytoplankton, for example, if adaptation to high temperatures would lead to a decrease in fitness at low temperatures, as previously observed in bacteria (Bennett & Lenski 1993) and bacteriophages (Knies *et al.* 2006).

The TRN for population growth is an emergent property of the temperature dependences of enzyme activities (Ratkowsky *et al.* 2005; Corkrey *et al.* 2014), and few processes or pathways within the cell should be immune to the effects of directional temperature selection (Nedwell 1999; Baker *et al.* 2016; Padfield *et al.* 2016). Depending on the genes affected (and thus the mechanism by which thermal adaptation is achieved), changes in maximum growth rate may be accompanied by changes in traits not directly associated with the TRN. For example, changes in stoichiometry in response to thermal adaptation (e.g. Schlüter *et al.* 2014) may be due to changes in relative resource requirements of phytoplankton cells, which in turn may affect species’ competitive abilities (e.g. for N and P) and ultimately global biogeochemical cycles (Redfield 1958; Litchman *et al.* 2007; Baker *et al.* 2016). An increase in maximum growth rate at the selection temperature may lead to a decrease in affinity for a given nutrient due to a tradeoff between allocation of cellular resources to reproduction versus nutrient uptake (Grover 1991; Klausmeier *et al.* 2004). If the nutrient in question is never limiting, selection may favor a high maximum growth rate over affinity, and the tradeoff may ultimately result in a weakening of competitive ability for that nutrient in the selection environment.

To address the need for studies of the broad physiological and ecological consequences of thermal adaptation in phytoplankton, we performed a long-term selection experiment investigating the effects of prolonged exposure to temperatures above and below the growth optimum on a suite of temperature-dependent traits in a model marine diatom, *Thalassiosira pseudonana*. We evolved this diatom in replicate populations at two different temperatures (16 and 31°C) for 18 months (~400 generations). Throughout this period, we monitored the maximum (nutrient-saturated) growth rates of the replicate populations (five at each temperature) and conducted temperature-dependent growth assays to determine whether and how thermal adaptation leads to evolutionary change in TRN for population growth and the traits associated with it: *T*_opt_, *CT*_max_, maximum growth rate (*μ*_max_) at *T*_opt_ (*μ*_opt_), *μ*_max_ at the selection temperatures, and the curvature of the TRN at and below *T*_opt_. We also conducted nutrient-dependent growth assays on all replicate populations at both selection temperatures to investigate the consequences of thermal adaptation for nitrate growth affinity, an important trait contributing to nitrate competitive ability (Tilman 1982)—i.e whether selection for allocation of resources to reproductive machinery over nitrate uptake machinery led to a trade-off between nitrate growth affinity and maximum growth rate (Grover 1997; Litchman & Klausmeier 2001).

## Methods

### Temperature selection experiment

We obtained a monoculture of *Thalassiosira pseudonana* CCMP1335 (Hustedt) Hasle et Heimdal from the Provasoli-Guillard National Center for Culture of Marine Phytoplankton (CCMP), Maine, USA, and isolated a single cell by plating on agarose ESAW marine culture medium (Harrison *et al.* 1980). From this progenitor, we propagated ten replicate populations (“selection lines” hereafter) into 20 ml ESAW marine culture medium contained in 50 ml Cellstar polystyrene tissue culture flasks with breathable caps (Greiner Bio-One GmbH, Frickenhausen, Germany), which we gently agitated daily to keep cells in suspension. We assigned five replicates to the 16°C treatment and the other five to 31°C. These two temperatures were below and above the previously-recorded thermal optimum (*T*_opt_) of *T. pseudonana* (~26°C: Boyd *et al.* 2013), chosen so that the maximum growth rates (*μ*_max_) at the two temperatures were roughly equal (~0.8 d^−1^ at the start of the experiment). Selection lines were maintained in temperature-controlled growth chambers under a 14:10 light:dark cycle, illuminated at 110 μmol photons m^−2^ s^−1^ during the day. All lines were maintained in ESAW medium for ~50 generations at a dilution rate of 0.5 d^−1^ (diluted daily), after which all cultures were transferred to L1 medium (Guillard & Hargraves 1993) due to poor culture health in ESAW. L1 was made using 43.465 g l^−1^ artificial sea salt (Tropic Marin, Wartenberg, Germany), which yields a specific gravity of 1.025, similar to natural seawater. Upon switching culture media, we also altered the dilution regime, such that 10^6^ cells (usually ~0.5-1.0 ml) were transferred to fresh medium every 4 d (with occasional deviations of ±1 d). The selection lines required an additional ~50 generations to adjust to the new dilution regime before consistent growth rates and culture health were achieved. These changes led to a substantial improvement in culture health and maximum growth rates. Cell densities of all replicate cultures were determined at each transfer for the remainder of the experiment using a CASY particle counter (Schärfe System GmbH, Reutlingen, Germany), and average population growth rates for each propagation period determined by taking log(*N*_t_*/N*_0_)/*t*, where *N* is the population density and *t* is time in days; growth assays in which cell densities were estimated daily were conducted periodically throughout the selection experiment, and did not at any point indicate that 4 d was long enough for populations to reach carrying capacity.

### Determination of temperature-dependent trait values

To evaluate evolutionary change in physiological traits, we conducted temperature-dependent (after ~350 generations) and nitrate-dependent (after ~400-450 generations) population growth assays on evolved selection lines. To derive temperature-dependent trait values, we fit a recent model proposed by Thomas *et al.* (2017), in which birth and death are both exponential functions of temperature. The resulting curve is a unimodal, left-skewed thermal reaction norm (TRN). The double-exponential model (DE model, hereafter) is as follows:

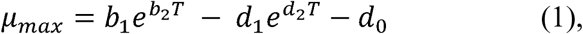

where *μ_max_* is the specific growth rate (d^−1^), *T*, *b_1_* and *d_1_* are the pre-exponential constants for birth and death, respectively, *b_2_* and *d_2_* are the exponential rates of increase in both terms, and *d_0_* is the temperature-independent death rate. We derived *T*_opt_ analytically by taking the derivative of the thermal performance curve between 10°C and 34°C and setting it equal to zero (see Appendix A). Per-capita population growth as a function of external nitrate concentration can be described using the Monod equation (Monod 1949):

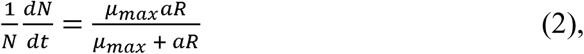

where *R* is the resource (nitrate, in this case), *μ*_max_ is the resource-saturated (maximum) population growth rate, and *a* is the nitrate affinity (the initial slope of the Monod curve). Note that this formulation directly incorporates the affinity *a*, rather than the traditional formulation with the half-saturation constant for growth, *K*. Affinity is a more useful parameter because it is more directly indicative of competitive ability (Healey 1980). The two parameters are related by *K* = *μ*_max_/*a*.

#### Temperature-dependent growth assays

For temperature-dependent growth assays we first sub-cultured all selection lines and acclimated sub-cultures to ten temperatures spanning the thermal niche for ~12 generations. Approximately 10^5^ cells from each acclimated culture were then propagated into fresh L1 medium, and each culture’s density estimated daily at the acclimated temperature until reaching stationary phase (5-10 d, depending on the temperature). Light conditions were as above. Culture density was estimated daily at all temperatures ≥16°C by placing the polystyrene culture flask in a Shimadzu UV-2401PC spectrophotometer (Shimadzu Corporation, Kyoto, Japan) and measuring Abs_436_ (the wavelength corresponding to the absorbance maximum of chlorophyll *a*) (Neori *et al.* 1986); Fogging of culture flasks at temperatures <16°C prevented use of this method, so daily density estimates at 3 and 10°C were conducted using the CASY counter. We calculated exponential growth rates by fitting regression lines to log-transformed population densities over time. Single cultures of each selection line were assayed at each temperature, except those at 16 and 31°C, which were assayed in triplicate to facilitate direct statistical comparison of growth rates. This comparison serves as a “reciprocal transplant” experiment, a common approach to test for local adaptation (Kawecki and Ebert 2004).

#### Nitrate-dependent growth assays

To determine if the nutrient-related traits changed as a result of adaptation to different temperatures, we estimated Monod growth curves for nitrate at 16 and 31°C. In particular, we sought to determine whether temperature selection had caused evolutionary divergence in nitrogen affinities between lines from each temperature treatment when assayed at both selection temperatures, and whether this divergence corresponded to a growth-affinity tradeoff. All ten selection lines were assayed in triplicate at each selection temperature and at ten N concentrations ranging from 0 to 882 μmol l^−1^ (the N concentration of undiluted L1 medium). Population growth rates were measured as above. Prior to the Monod assay, all assayed lines were acclimated to assay temperatures as above, followed by a 5 day N-starvation period in which cultures were kept in modified N-free L1 medium.

#### Model fitting, trait estimation and statistical analysis

We conducted all statistical analyses using R statistical programming language, version 3.3.2. We fit Equation (1) to temperature-dependent growth data using the “mle2()” function from the “bbmle” package (Bolker 2016). To fit Equation (2) to nutrient-dependent growth data, we used weighted, nonlinear least squares regression, due to high uncertainty in some growth rate estimated for these assays; these analyses were carried out using the “gnls” function from the “nlme” package (Pinheiro *et al.* 2017), with a fixed variance function (“varFixed()”). Estimation of uncertainty in parameter estimates was identical to the nonparametric, residual bootstrapping procedure described in Listmann *et al.* (2016).

We compared bootstrapped TRN trait distributions and statistically fit means and variances of nitrate affinity using simple linear models with selection line as the only predictor (see Supplemental materials section S1.1). We compared *μ*_max_ and nitrate affinities at the selection temperatures within and among selection lines (“reciprocal transplant” approach) using linear models with selection line and assay temperature as predictors (see Supplemental materials section S1.2). We did not fit an interaction term due to insufficient degrees of freedom. Bootstrapped nitrate affinity (*a*_NO3_) estimates were log-normally distributed and were log-transformed for statistical fits and comparisons (i.e. all affinity estimates were entered as exp[ln(*a*)] for statistical fits, and 95% CI are for ln(*a*)).

## Results

### Evolutionary change in thermal reaction norms

The growth for 350 generations at two different temperatures, below and above the temperature optimum for *T. pseudonana* (16 and 31°C), led to a significant divergence of the thermal reaction norms (Figure 1). *T*_opt_ was 2.1°C higher, on average, in selection lines evolved at 31°C than in those evolved at 16°C (Figure 2A; see standardized linear model effects with 95% CI for individual replicates in Supplemental Materials Figure S1), as was the maximum growth rate at *T*_opt_ (*μ*_opt_) (Figure 2B; SM Figure S1). *CT*_max_ was also higher in 31°C-selected lines than in 16°C-selected lines, with one exception (replicate line 31.5). However, divergence in *CT*_max_ was smaller than for *T*_opt_, <1°C on average (Figure 2C; SM Figure S1).

**Figure 1.**
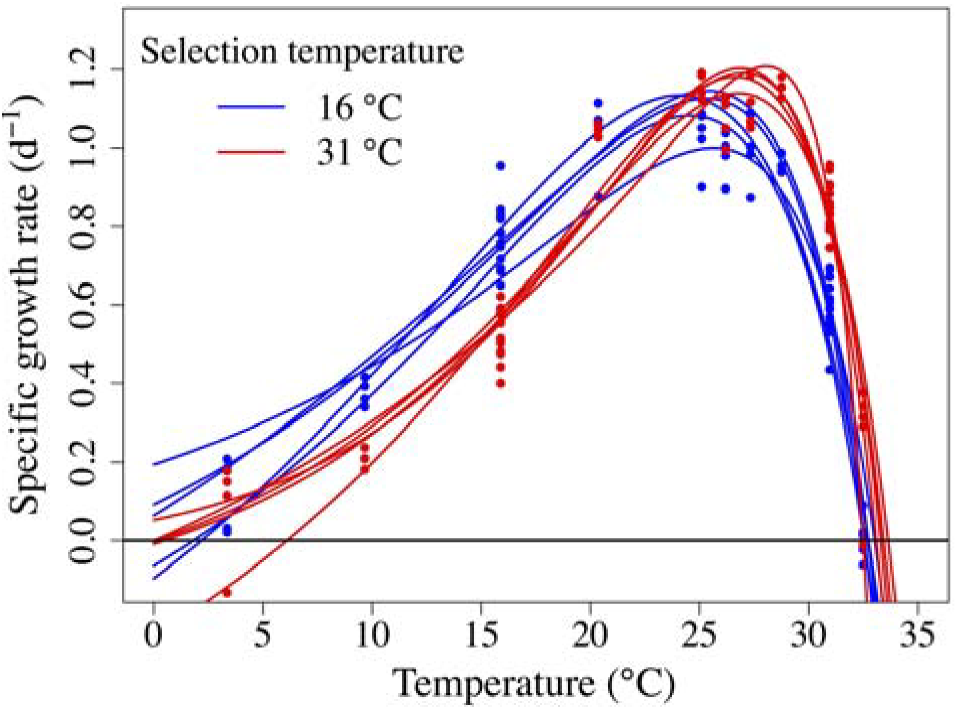
Thermal reaction norms for per-capita population growth of 10 *T. pseudonana* populations after 350 generations of experimental selection, five at 16°C (blue) and five at 31°C (red). Curves were fit using maximum likelihood estimation.

**Figure 2.**
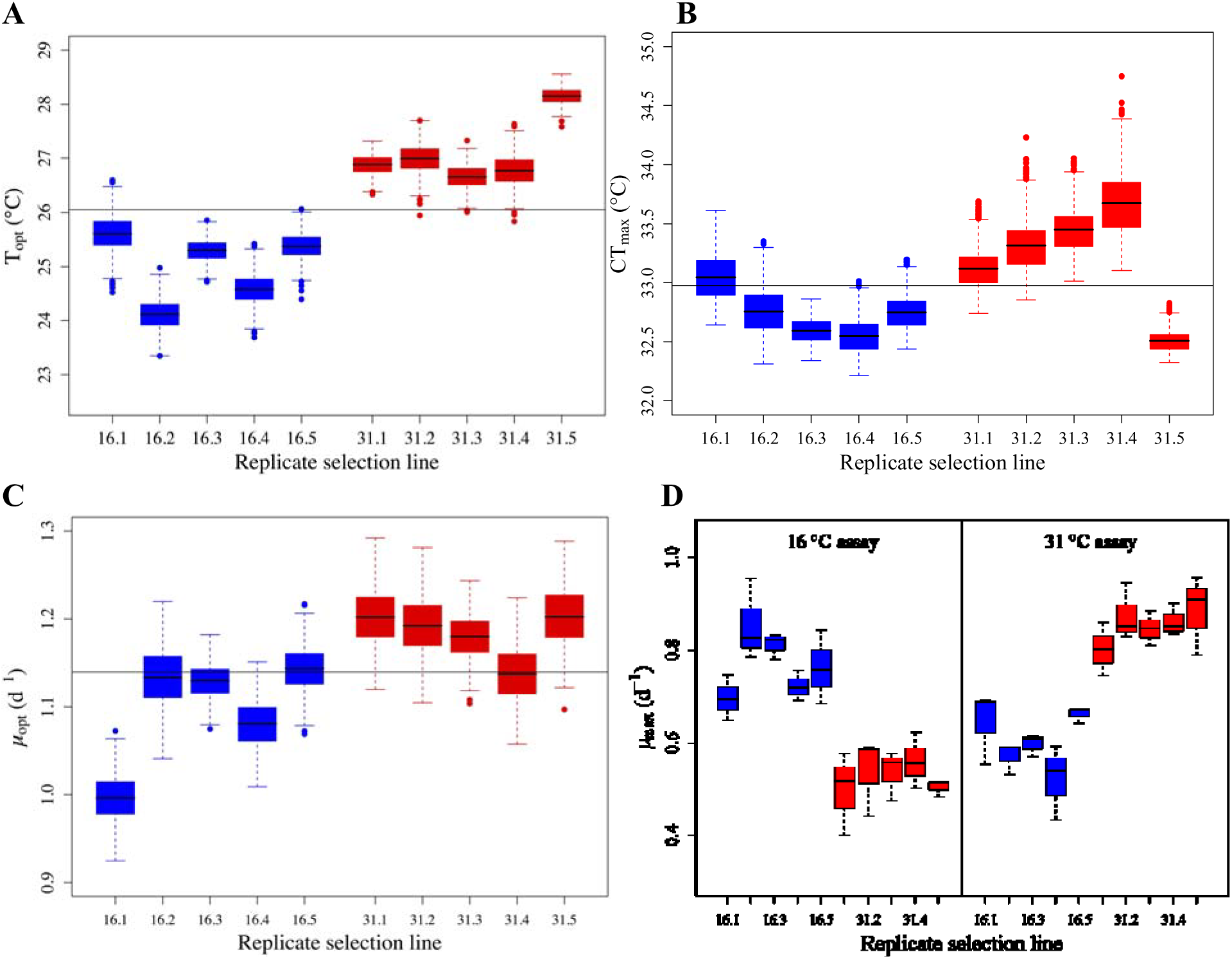
(A) Box-and-whisker plots of bootstrap *T*_opt_ estimates (n = 1000). Boxes span the first-third quartiles; dots are outliers; black lines inside boxes represent *T*_opt_ estimates from the DE fits in panel A. (B) Box-and-whisker plots of *CT*_max_ bootstrap distributions. Symbols are as in panel B. (C) Box-and-whisker plots of *μ*_max_ bootstrap distributions. Symbols are as in panel B. (D) Box-and-whisker plots of *μ*_max_ in the reciprocal transplant scenario (both selection groups assayed at both selection temperatures). These assays were conducted in triplicate (no bootstrapping). In panel D, black horizontal lines are median values for each selection line. In panels B and C, the thin, horizontal line across the whole panel is the grand mean across all bootstrap parameter values, to aid in visualization of trait differences among selection lines.

A reciprocal transplant comparison between temperature treatments revealed that all selection lines from each treatment had higher maximum growth rates (*μ*_max_) in their “home” temperature environments than in “away” environments, with the exception of replicate line 16.1; “local” lines also had higher growth rates than “foreign” lines in all cases (Figure 2D; SM Figure S1). Taken together, these results are the signature of local adaptation to temperature (Kawecki & Ebert 2004; Blanquart *et al.* 2013). While *μ*_max_ estimates differed slightly between 350 and 450 generations (measured for TRN and Monod assays, respectively), differences were not directionally consistent across selection lines, and the overall pattern of local adaptation persisted (Figure 5A).

The peaks of the TRN of 31°C-selected lines appeared sharper, and the left tails more deeply concave-up than those of 16°C-selected lines, both for the parametric (Fig. 1) and nonparametric (Fig. S5) TRN. These observations, combined with the extremely narrow range of *CT*_max_ between the two selection regimes, led us to hypothesize that *CT*_max_ is less evolutionarily labile than *T*_opt_, resulting in greater skew in TRN of 31°C-selected lines, rather than a simple rightward shift of the TRN along the temperature axis. The difficulty in reliably estimating *CT*_min_ prevented statistical comparisons of *CT*_min_ and of thermal niche width among selection lines, and also prevented estimation of skewness. However, we determined the distance between *T*_opt_ and *CT*_max_ for all bootstrapped thermal reaction norms; ideally, this distance would be scaled by thermal niche width, but absence of reliable thermal niche width estimates prevented scaling. The distance between *T*_opt_ and *CT*_max_ (“upper tail length”) was smaller in 31°C-selected lines than in 16°C-selected lines (Figure 3A; SM Figure S2), suggesting that the change in *T*_opt_ was greater than the change in *CT*_max_ at one or possibly both selection temperatures. We also determined the “sharpness” of the TRN peaks by taking the second derivative of the TRN with respect to temperature at *T*_opt_ (“Peak sharpness” [Figure 3B] is the negative of this number), and the upward-concavity below *T*_opt_ by comparing the maxima in the second derivative across 0 ≤ *T* ≤ *T*_opt_. We estimated the inflection point of each TRN by setting 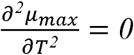 and solving for T within the range 0 ≤ *T* ≤ *T*_opt_. 31 °C-selected lines had sharper peaks at Topt, were more deeply concave-up below Topt, and had inflection points at higher temperatures than 16°C-selected lines (Figure 1; Figure 3B-3D; SM Figure S2), suggesting greater negative skewness. The area under each TRN between 0°C and *CT*_max_ (°C d^−1^), estimated using the “auc()” function in the “flux” R package (Jurasinski et al. 2014), was smaller in 31 °C lines than in 16°C lines in four out of five cases; AUC for replicate line 16.4 was comparable to those for 31°C-selected lines (Figure 3E; Figure S2).

**Figure 3.**
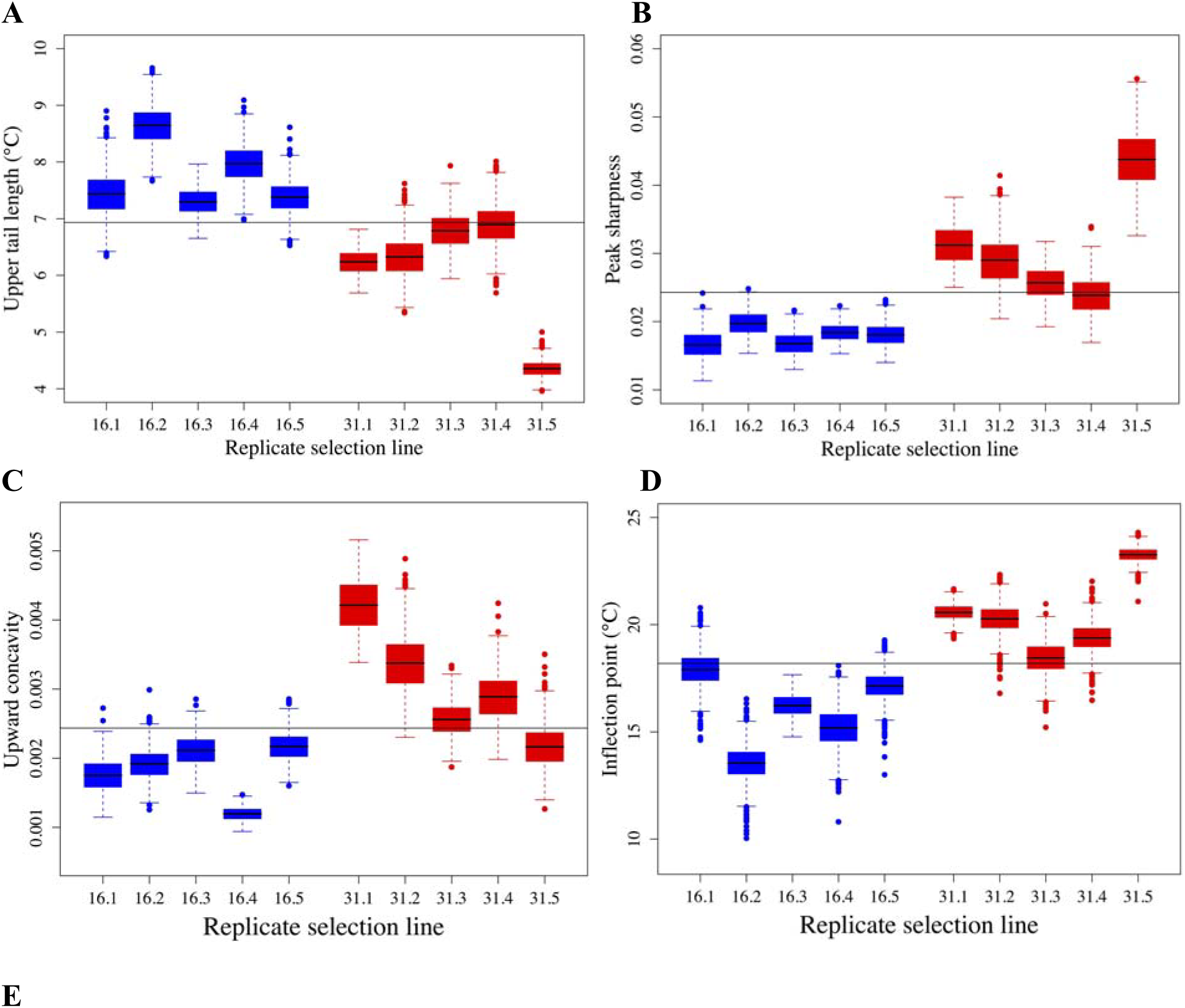
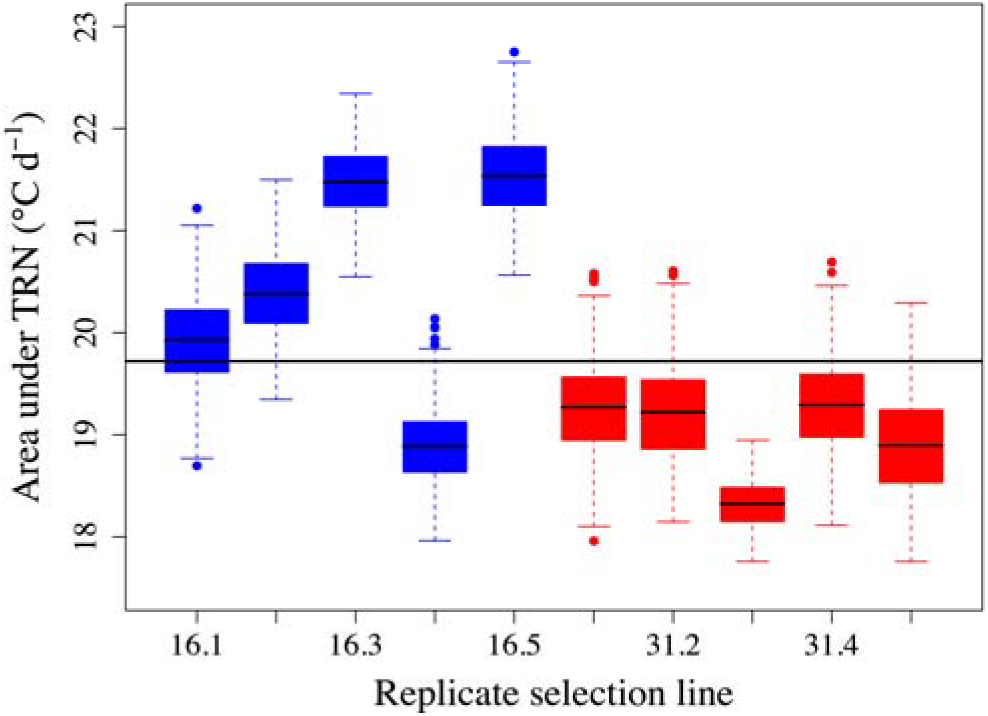
(A) Upper tail lengths (*CT*_max_ − *T*_opt_) derived numerically from bootstrap DE curves (n = 1000). (B) Peak sharpness of bootstrap DE curves. Sharpness is calculated as the negative of the second derivative of the DE curve with respect to temperature at *T*_opt_ (larger values represent more negative gradients). (C) Upward concavity of bootstrap DE curves below *T*_opt_, calculated as the maximum of the second derivative across 0 ≤ *T* ≤ *T*_opt_. (D) The inflection point below *T*_opt_, where 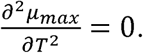 All symbols are as in Figure 1 B,C. (E) Area under the TRN (AUC) (°C d^−1^), determined by taking the integral the fit TRN between 0°C and *CT*_max_.

Adaptive divergence in *μ*_max_ at selection temperatures was likely driven mostly by evolution of 31°C-selected lines. Maximum growth rates in the 16°C-selected lines first increased between 150 and 350 generations, then declined between 350 and 500 generations, yielding a net change near zero. 31°C-selected lines, however, first increased *μ*_max_ rapidly for ~300 generations, after which little change occurred (Figure S3). Between 450 and 500 generations, all five 31°C-selected lines again appear to increase *μ*_max_ (Figure S3), though drawing inferences from this short timeframe may be premature.

### Evolution of competitive ability for nitrate

Temperature selection led to evolutionary change in traits beyond the thermal reaction norm for growth rate. Specifically, nitrate growth affinities, derived from nitrate-dependent growth (Monod) assays (Figure 4), diverged between temperature treatments, leading to changes in competitive ability for nitrate in both “home” and “away” environments. While all selection lines remained locally adapted to their selection temperatures in terms of their maximum growth rates (*μ*_max_) after 450 generations (Figure 5A), variation in nitrate affinity (*a*_NO3_) was more consistent between assay temperatures than between selection groups; *a*_NO3_ was higher for all but three (replicate lines 16.1, 31.1, 31.5) when assayed at 31°C, regardless of selection temperature (Figure 5B). Three out of the five 31°C-selected lines (replicate lines 31.2, 31.3, 31.5) had marginally higher *a*_NO3_ than all of the 16°C-selected lines when assayed at 16°C, with one (31.1) dramatically higher (Figure 5B). However, at 31°C, two 16°C-selected lines (16.3, 16.4) and two 31°C-selected lines (31.2, 31.3) had *a*_NO3_ > 0.05 l μmol^−1^ d^−1^, while the rest were nearly indistinguishable, with affinities around 0.03 l μmol^−1^ d^−1^ (Figure 5B). The trade-off between the maximum growth rate and nitrate affinity was apparent in the 16°C-selected lines but, interestingly, was weak or absent in the 31°C-selected lines, with replicates pooled across assay temperatures (Fig. S4).

**Figure 4.**
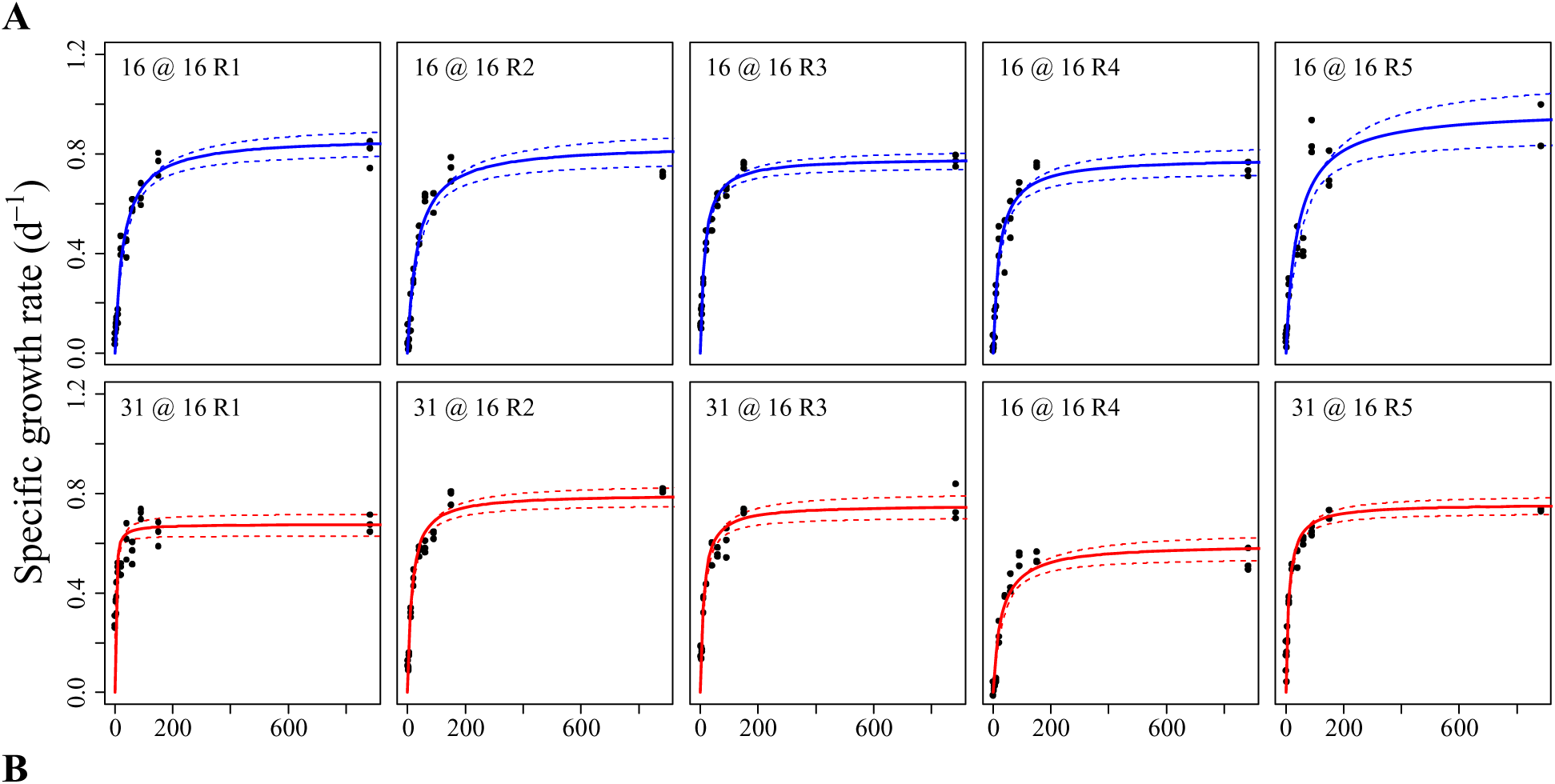
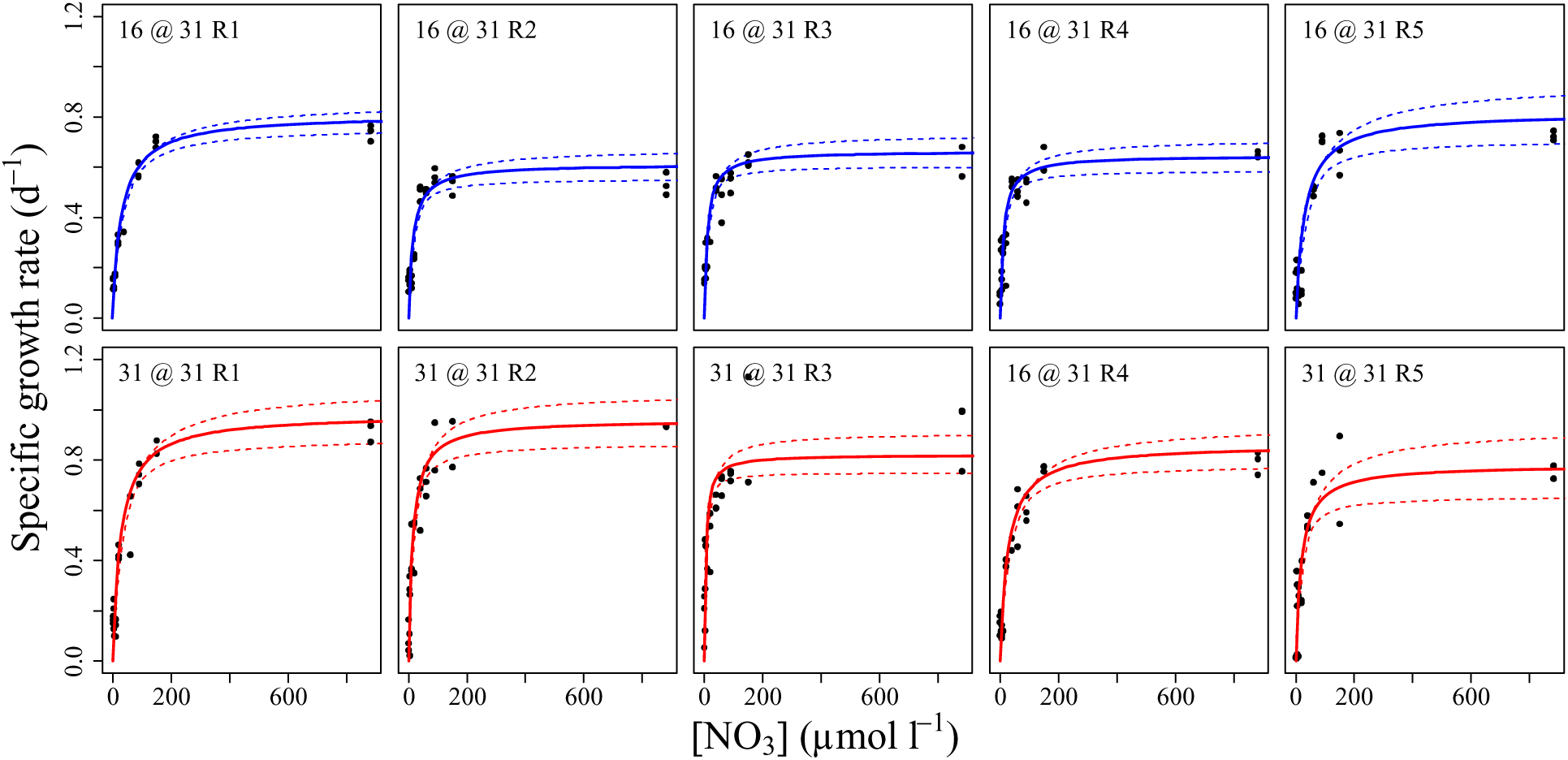
Nitrate-dependent growth (Monod) assays at 16°C (A) and 31°C (B). Curves were fit to *T. pseudonana* exponential growth rates at ambient NO_3_ concentrations ranging from 0 to 882 μmol l^−1^ (the NO_3_ concentration in unaltered L1 marine medium). Monod parameters (*a*_NO3_ and *μ*_max_) were estimated using generalized leased squares.

**Figure 5.**
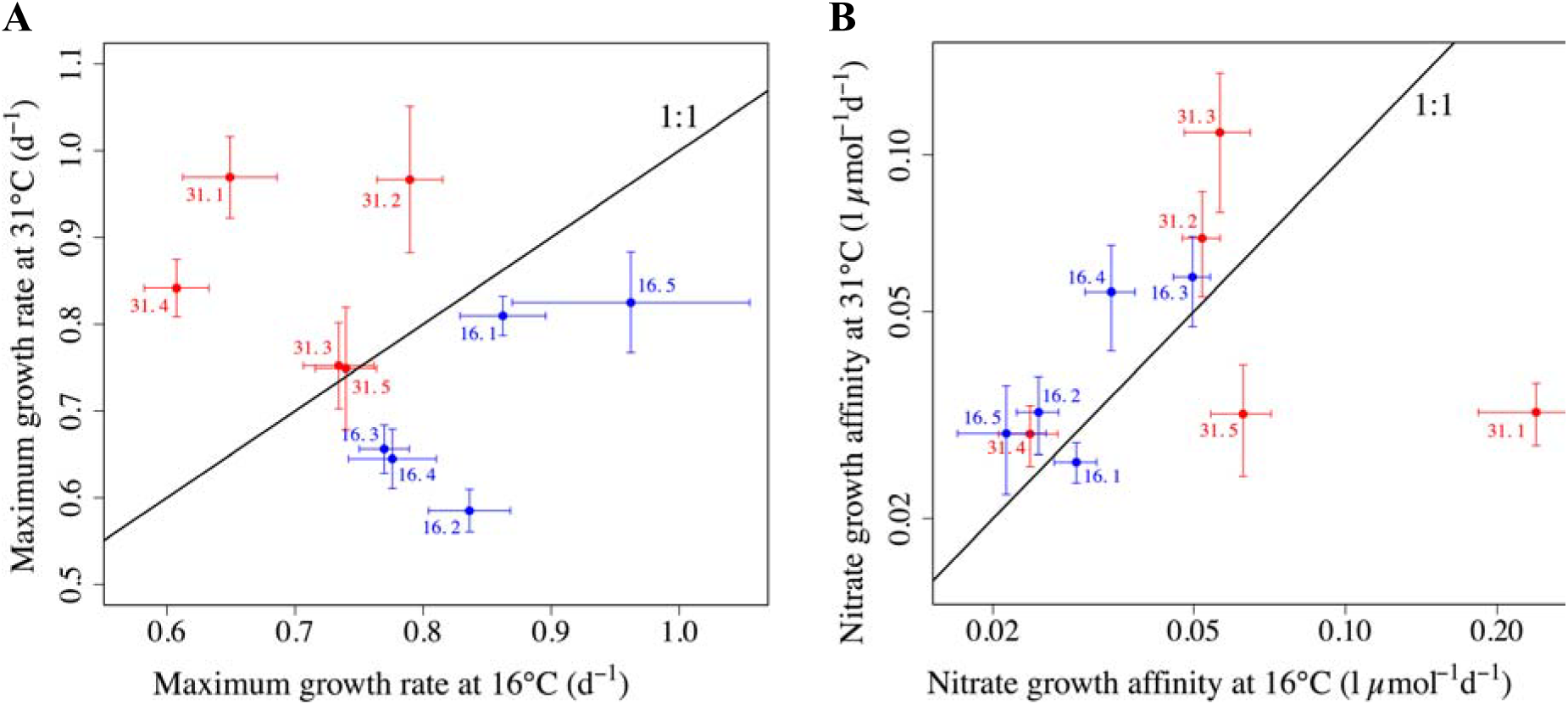
Nitrate-dependent growth (Monod) kinetic parameters. (A) *T. pseudonana* maximum specific growth rates (*μ*_max_) for 31°C- and 16°C-selected lines (red and blue, respectively) assayed at 31°C versus at 16°C. (B) Nitrate growth affinity (*a*_NO3_); colors and assay temperatures are as in panel A. Note that the y-axis in panel B is on a log scale. Black diagonals are 1:1 lines. Bars are ±1 SE.

## Discussion

Evidence that phytoplankton can evolve rapidly in response to environmental change is mounting (Collins & Bell 2004; Hutchins *et al.* 2015; Listmann *et al.* 2016), but the response to a single, directional selection pressure can be multifaceted (Low-Décarie *et al.* 2013; Schlüter *et al.* 2014; Hutchins *et al.* 2015). We showed that adaptation to different temperatures not only changes the growth rate at the selection temperature but significantly alters the shape of the whole thermal reaction norm (TRN), a function-valued trait. The changes included shifts of critical temperature traits (e.g. *T*_opt_ and *CT*_max_), changes in the slope and curvature of the TRN at, below, and above *T*_opt_, and changes in the total area under the TRN (“AUC”, hereafter). Adaptation to high temperature resulted in a growth rate decline at low temperatures (a performance trade-off). Moreover, traits not directly associated with the TRN for population growth, such as competitive abilities for nutrients, changed in response to temperature selection as well, possibly due to pleiotropic effects (Elena & Lenski 2003) or resource allocation or acquisition tradeoffs (Gilchrist 1995; Angilletta *et al.* 2003).

Selection at 31°C, well above the previously recorded *T*_opt_ for *T. pseudonana* (Boyd *et al.* 2013), led to a shift in *T*_opt_ of ~2 °C, on average. Concurrent with this shift, the maximum growth rate at *T*_opt_ (*μ*_opt_) increased by ~0.1 d^−1^ (“vertical shift”: Izem & Kingsolver 2005) and all strains but one (replicate line 16.1) met both the “local versus foreign” and “home versus away” criteria for demonstrating local adaptation at both 16°C and 31°C (Kawecki & Ebert 2004; Blanquart *et al.* 2013). We are unsure what caused the decline in growth rate of the 16°C-selected lines after ~350 generations (Figure S3); a reduction in growth rates in evolution experiments was observed previously and could also be attributed to the accumulating cellular damage associated with the initial evolution of faster growth rate (“Prodigal Son” dynamics) (Collins 2016). A second possible explanation is clonal interference, though determination of relative clone frequencies within a population would require identification and quantification of genetic markers (Gerrish & Lenski 1998). Despite this reduction in fitness, however, local adaptation was still apparent in 16°C-selected lines at 450 generations (Figure 5).

The results presented here suggest that the variation in *T. pseudonana*’s TRN caused by thermal adaptation led to (or was driven by) a number of trade-offs. First, while we could not precisely estimate *CT*_min_ or the thermal niche width, the performance trade-off resulting from selection at 16°C versus at 31°C, combined with changes in *T*_opt_, strongly suggest some horizontal shift in the TRN (Izem & Kingsolver 2005; Kingsolver *et al.* 2009), as previously observed in bacteriophages (Knies *et al.* 2006) and the marine coccolithophore *Emiliana huxleyi* (Listmann *et al.* 2016). In addition, both temperature “generalist-specialist” and resource allocation trade-offs were apparent (Gilchrist 1995; Angilletta *et al.* 2003). The higher, sharper peaks and compressed upper tails in the TRN of 31°C-selected lines, accompanied by a reduction in fitness at 16°C, suggest greater specialization for growth at high temperatures in 31°C-selected lines and less specialization in general in 16°C-selected lines. However, the reduction in area under the TRN curve (“AUC”) in 31°C-selected lines suggests the existence of an acquisition or allocation trade-off as well. Gilchrist (1995) predicted that, absent a major shift in resource acquisition or allocation toward or away from reproduction, the AUC should remain constant—in other words, the derivative of the function may change, but its integral does not (Gilchrist 1995; Angilletta *et al.* 2003). We found that, assuming a hard cutoff in positive population growth at 0°C, 31°C-selected lines had a lower total AUC than four out of five 16°C-selected lines (Figure 3E), despite having higher maximum growth rates at and above *T*_opt_.

We propose a modification to Equation (1) that leads to a prediction of greater temperature specialization in high-temperature adapted strains while also allowing for an allocation trade-off in which enhanced enzyme repair machinery at high temperatures comes at a cost to reproduction at temperatures below *T*_opt_. In high-temperature adapted species, the thermodynamically predicted temperature of maximum enzyme stability falls well below the statistically fit *T*_opt_ (Ratkowsky *et al.* 2005; Corkrey *et al.* 2014), suggesting that it is rapid repair, rather than enhanced stability of enzymes that allows for positive growth at very high temperatures. The trade-off inherent in the following model simply assumes a lack of plasticity in resource allocation to enzyme repair at high temperatures versus reproduction across temperatures (Angiletta *et al.* 2003).

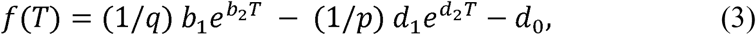

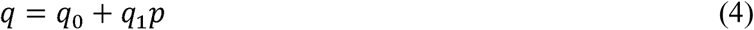

Here, the exponential mortality term is weighted inversely by *p*, which represents investment in protection from heat-induced denaturation of enzymes (or the induction of repair enzyme activity); *q* is the resource content of a cell (quota), which is linearly related to *p*. Thus, as investment in protection and repair increases, temperature-dependent mortality decreases. However, temperature-dependent birth is also negatively affected as resources are diverted from reproduction and allocated to protection. The modified DE model can produce changes in the curvature of the TRN similar to those observed in the *T. pseudonana* selection experiment (Figure 6). The model predicts that peak sharpness at *T*_opt_, upward concavity below *T*_opt_, and the location of the inflection point are all positive, saturating functions of *p* (Figure S12A-D).

**Figure 6.**
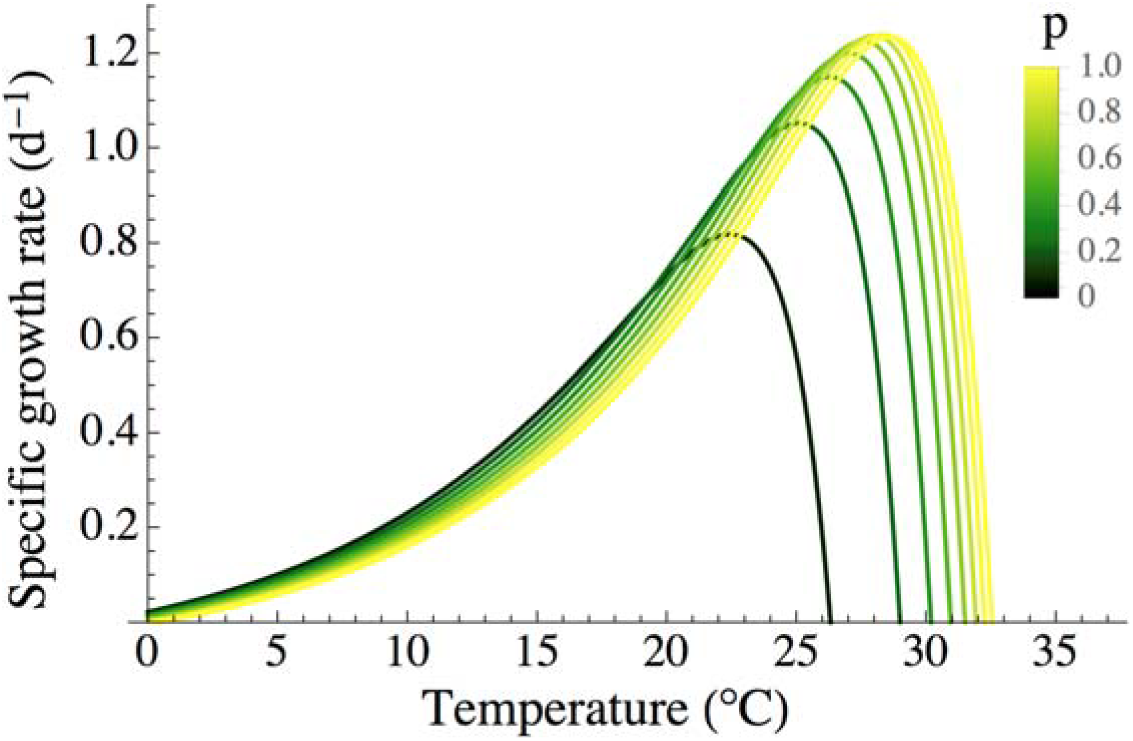
The modified double-exponential model with protection (p) ranging from 0.1 to 1. Other parameter values are as follows: b_1_ = 0.124, b_2_ = 0.1, d_0_ = 0.1, d_1_ = 3.059×10^−7^, d_2_ = 0.5.

There are two observed results that this model does not predict, however. First, the model predicts that upper tail length is a positive, saturating function of *p* (Figure S7). In contrast, the upper tails of our experimentally derived thermal reaction norms were more compressed in 31°C-selected lines, compared to 16°C-selected lines (Figure 3A). Second, in experiments, the change in *CT*_max_ was smaller than that in *T*_opt_—these two discrepancies are likely not independent of one another. Evolution of *CT*_max_ may thus be constrained by evolutionary barriers not incorporated in the model. Although evolution of *CT*_max_ in response to temperature selection has been observed in the bacterium *Escherichia coli* (Mongold *et al.* 1996) and in the marine coccolithophore *Emiliana huxleyi* (Listmann *et al.* 2016), the small changes in *CT*_max_ relative to *T*_opt_ in this study suggest that, at least in *T. pseudonana*, *T*_opt_ is more evolutionarily labile than *CT*_max_, as suggested by Araújo *et al.* (2013)—although the mere existence of vast diversity in *CT*_max_ in nature (see, e.g., Corkrey *et al.* 2014) indicates that such constraints are not evolutionarily insurmountable. However, on timescales relevant to the current rate of increase of global sea surface temperature, constraints on evolution of a higher *CT*_max_ may cause regional extinctions, contributing to diversity loss, especially at low latitudes (Thomas *et al.* 2012).

An increase in the maximum growth rate (fitness) is the most predictable response to the temperature selection environment, but the physiological causes and ecological consequences of that change can be indirect and far-reaching (Schlüter *et al.* 2014; Padfield *et al.* 2016). In the presence of a growth rate-competitive ability tradeoff, for example, selection for rapid growth (high *μ*_max_) in nutrient-replete conditions (as was the case here) may lead to a decline in competitive ability (suggested here by a decline in *a*_NO3_), though as we observed, this tradeoff may be escapable. Whether or not a tradeoff is observed may depend on the affected genes and the mechanisms by which a population achieves thermal adaptation. Given the diversity in *a*_NO3_ and among selection lines within a single selection and assay temperature group, adaptation to temperature was possibly driven by changes to a distinct combination of loci in each selection line—mutations in some lines were relevant to NO_3_ uptake and metabolism (e.g. genes for cell membrane NO_3_ transporters, NO_3_ reductases and plastid-localized nitrite transporters) (Armbrust *et al.* 2004), while others may have affected other systems, such as photosynthesis or reproductive machinery (Padfield et al. 2016).

In the absence of nutrient limitation (and thus any selection for enhanced competitive ability), selection likely favors high *μ*_max_ over high nitrate affinity. However, thermal adaptation in the ocean would occur under nitrogen limitation in most cases, at least in the temperate ocean where *T. pseudonana* is commonly found (Fong 2008), and may thus produce competitive abilities quite different from those we observed here. While we acknowledge the challenges inherent in combined temperature-nutrient selection experiments (e.g. longer generation times and high mortality due to failure to meet minimum nitrate quotas), we suggest that future temperature selection experiments must account for nutrient limitation to reflect realistic climate change scenarios in natural systems.

Evolution experiments are essential to enhancing our understanding of the effects of climate change on marine and freshwater food webs. Arguably, phytoplankton are among the highest-payoff organisms upon which we can conduct evolution experiments; they offer a unique combination of potential for rapid evolutionary responses to environmental change, tractability as experimental subjects, and global ecological importance (Reusch & Boyd 2013). Eco-evolutionary responses to changes in temperature, ocean acidity, nutrient limitation and other environmental factors are complex, and are made more complex by interactions among these factors. Single-stressor evolution experiments are a valuable first step, but future evolution experiments must account for interactive effects of multiple stressors (e.g. temperature and nitrate limitation).

## Acknowledgements

We thank Colin Kremer for statistical consultation and Mridul Thomas for culturing tips and conceptual input. This work would have been impossible without the support of our technicians at Kellogg Biological Station, Allyson Hutchins and Pamela Woodruff. DRO was supported by the National Science Foundation Graduate Research Fellowship, and by the Kellogg Biological Station Undergraduate Mentor Fellowship. This work was funded by National Science Foundation Grants OCE 0928819 and 1638958 to EL and CAK.

